# Proteome-Scale Mining and Multi-Objective Prioritization of Encrypted Antimicrobial Peptides with Experimental Validation

**DOI:** 10.64898/2026.08.18.745516

**Authors:** Qingxiu Li, Zhenjun Li

## Abstract

Encrypted antimicrobial peptides (eAMPs) are bioactive fragments embedded within larger proteins and represent an underexplored source of antimicrobial candidates. We developed a multi-layer proteome-mining framework to identify and prioritise eAMPs from 95%-identity-reduced protein sets derived from 265 high-quality bacterial genomes. Three complementary, layer-specific extraction strategies targeting protein termini, internal cleavage sites, and cationic hotspots yielded 29,251,180 unique peptide candidates. Dual AMP prediction with AMP-scanner v2 and Macrel reduced this space to 3,249,772 consensus candidates. Downstream prioritisation followed two complementary routes: a low-haemolysis branch focused on selectivity-oriented candidates and a high-activity branch that retained predicted haemolytic sequences as mechanistic comparators. Structure prediction and review were performed for 185 candidates, and 18 entered Tier-1 developability, novelty, and membrane-activity assessment. Three sequence-matched representatives were selected for experimental evaluation. Molecular-dynamics simulations supported water-phase stability of GEAMP_71c139393ac596b5 and deep anionic-membrane insertion by GEAMP_12ffb5d589c8cb1b. In replicated colony-count assays against Escherichia coli and Staphylococcus aureus, all three peptides showed concentration-dependent activity over 8 – 128 μM. GEAMP_12ffb5d589c8cb1b was the most active, producing 1.52- and 2.27-log10 reductions, respectively, at 128 μM relative to the matched 8 μM condition. Together, these results establish a sequence-traceable workflow linking proteome-scale eAMP discovery with structural prioritisation and experimental activity assessment.

## Introduction

Antimicrobial resistance is a major and growing threat to global health, motivating the development of anti-infective modalities that complement conventional small-molecule antibiotics [1,14–17]. Antimicrobial peptides (AMPs) are attractive in this context because short cationic and amphipathic sequences can interact rapidly with bacterial envelopes, while the accessible sequence space of short peptides is exceptionally broad [2,3,15,16,18]. However, a high predicted probability of antimicrobial activity is insufficient for candidate selection. The same physicochemical properties that favour membrane interaction, including positive charge and hydrophobicity, can also increase erythrocyte lysis, mammalian-cell toxicity, aggregation, and poor developability. A useful discovery framework must therefore integrate activity-related predictions with safety, structural plausibility, novelty, and experimental feasibility.

Encrypted peptides provide one route to expand this discovery space. These peptides are short bioactive segments embedded within larger proteins and may become releasable through proteolytic processing [4,5,19–22]. Mining long proteins for encrypted AMP-like segments enables exploration beyond annotated AMP precursor families, but it also imposes an important interpretative boundary: an in silico extracted sequence is a hypothesis about a potentially releasable bioactive fragment, not direct evidence that the peptide is produced in vivo, adopts the predicted conformation, or selectively kills bacteria.

Machine-learning methods have extended AMP discovery from curated peptide collections to genome- and microbiome-scale datasets [6,7,10,11,23–32]. Because individual classifiers can inherit biases from their training data, we designed a staged workflow in which complementary extraction rules are followed by two-model AMP prediction, predicted haemolysis filtering, explicit multi-objective scoring, structural review, focused developability assessment, novelty analysis, and candidate-specific molecular-dynamics (MD) evidence. This design keeps distinct evidence classes separate rather than collapsing them into a single unqualified lead score.

Here, we apply this framework to nr95 protein collections from 265 high-quality bacterial genomes. The analysis begins with three mechanistically distinct eAMP-mining layers, proceeds through dual-model AMP consensus and two complementary downstream prioritisation routes, and then selects three sequence-defined representatives for focused structural, MD, and colony-count evaluation. Importantly, the same GEAMP identifiers and peptide sequences are carried from computational prioritisation into experimental testing, enabling direct sequence-level integration of the discovery and evaluation stages.

## Results

### Three complementary extraction layers generated a large but traceable eAMP candidate space

Proteins from 265 high-quality bacterial genome-derived nr95 collections were filtered to retain parent proteins of at least 100 amino acids and were scanned by three complementary extraction routes (Fig. 1). Layer 1 (L1) sampled N- and C-terminal regions, Layer 2 (L2) sampled neighbourhoods around basic cleavage-like motifs, and Layer 3 (L3) identified internal cationic hotspots followed by local fine-scale extraction. The three layers did not use a common scoring function. L1 used an eight-point relative peptide score (RPS) integrating terminal proximity, peptide length, cleavage context, and net positive charge and retained candidates with RPS ≥7. L2 used an independent eight-point cleavage-centred score and required the maximum score of 8, whereas L3 used an independent eight-point charge/length/hydrophobicity score and likewise required a score of 8.

**Figure 1.**
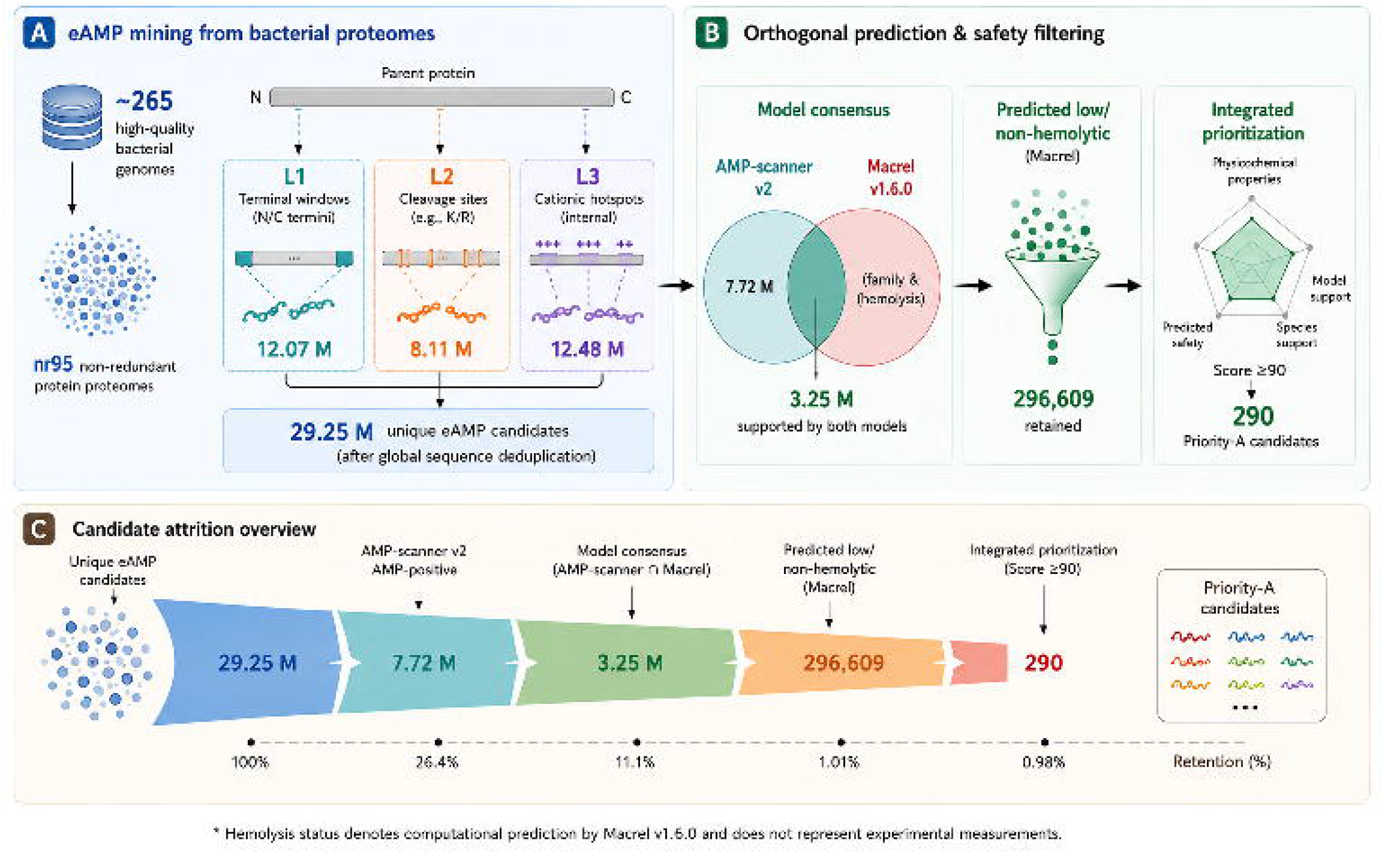
Multi-layer workflow for encrypted antimicrobial peptide discovery and prioritisation. Parent proteins (≥100 aa) from 265 nr95 bacterial proteome collections were interrogated by terminal scanning (L1), cleavage-neighbourhood extraction (L2), and internal cationic-hotspot mining (L3). The layers used distinct eight-point extraction scores: L1 RPS ≥7, L2 score=8, and L3 score=8. After exact sequence deduplication and AMP-scanner/Macrel consensus prediction, structural nomination proceeded through two parallel routes. The low-haemolysis route applied predicted safety filtering and multi-objective prioritisation, whereas the high-activity route ranked the same dual-model consensus without haemolysis exclusion and retained haemolysis predictions as risk annotations. The two routes nominated 32 and 153 structural candidates, respectively. Numbers indicate retained candidates and final structural-review dispositions.

L1, L2, and L3 generated 12,070,288, 8,111,517, and 12,483,768 sequence records, respectively. Exact amino-acid-sequence deduplication across all three routes produced 29,251,180 unique peptide candidates. Multi-route discovery was retained as provenance rather than discarded, allowing a candidate to carry evidence from more than one extraction logic. The three routes therefore broadened sequence sampling while preserving the biological hypothesis under which each fragment was recovered.

### Dual-model consensus supported distinct safety-oriented and high-activity prioritisation routes

The 29,251,180 unique candidates were first screened with AMP-scanner v2, which retained 7,715,263 sequences (26.4%). Requiring concordant AMP classification by Macrel v1.6.0 retained 3,249,772 candidates (42.1% of the AMP-scanner-positive pool). For the safety-oriented route, predicted low/non-haemolysis status, defined by a Macrel NonHemo label or a haemolysis probability below 0.5 in the haemolysis-prediction layer, reduced this consensus set to 296,609 sequences (9.1% of the dual-model pool).

The safety-oriented pool was scored on a 100-point scale combining AMP-related physicochemical properties (40 points), model evidence (30 points), cross-species conservation (20 points), and safety evidence (10 points). This produced 290 Priority A candidates (score ≥90), 177,457 Priority B candidates (80–89), 115,447 Priority C candidates (70–79), and 3,415 Priority D candidates (<70). These priority labels were used for candidate allocation within the safety-oriented route and were not interpreted as calibrated probabilities of biological activity or clinical success.

### Structure prediction compared two independently prioritised candidate routes

Structural candidate nomination was performed in parallel from the 3,249,772-member dual-model consensus. The low-haemolysis route applied the safety-oriented filtering and prioritisation described above and nominated 32 candidates for structure prediction. In parallel, the high-activity route did not exclude candidates on the basis of predicted haemolysis; haemolysis status was retained only as a risk annotation, and candidates were ranked by the predefined HighActivityScore, yielding 153 structural candidates. The top-200 ranking sets from the two route-specific prioritisation schemes showed no overlap, indicating that the routes sampled distinct regions of candidate space.

A total of 185 candidates were modelled with OmegaFold and ColabFold and underwent standardised structural review. In the low-haemolysis route, 15 candidates were classified Final_A, 13 Final_B, three Review, and one Exclude. In the high-activity route, 92 candidates were classified Final_A and 61 Final_B, with no Review or Exclude assignments. Across both routes, this yielded 107 Final_A, 74 Final_B, three Review, and one Exclude candidates. Final_A required pLDDT ≥75, predicted α-helical fraction ≥0.50, image-QC pass, and Priority A source status; Final_B required pLDDT ≥65 and predicted α-helical fraction ≥0.35. Across Final_A candidates, the mean pLDDT was 82.9, the mean predicted helical fraction was 0.71, and the mean net charge was +6.9.

#### Tier-1 assessment integrated sequence liability, developability, membrane activity, and novelty

Eighteen candidates entered Tier-1 assessment. Sequence-liability review classified 17 candidates as low risk and one as medium risk, with no high-risk candidates. Eleven candidate pairs showed >85% sequence similarity, highlighting redundancy that was considered during final representative selection. Aggregation-related assessment identified no high-aggregation-risk candidates; the mean amphipathicity score was 10.0. Synthesis feasibility was classified as easy for 17 candidates and moderate for one. Membrane-activity prediction classified 12 candidates as strong and six as moderate. The final Tier-1 set was divided evenly into nine Tier1_A and nine Tier1_B candidates.

Candidate annotation and novelty assessment combined functional annotation, database similarity searches, sequence clustering, and motif inspection. Novelty labels were used as a prioritisation feature rather than proof that a peptide represents a previously unrecognised biological family. Short water-phase OpenMM simulations and the availability of membrane-oriented simulation evidence were incorporated as additional candidate-specific evidence rather than mandatory filters for all 18 candidates.

### Three final candidates represented complementary selectivity and membrane-activity hypotheses

Three sequence-defined representatives were selected for focused analysis (Fig. 2; Table 3). GEAMP_71c139393ac596b5 (eAMP-01; GLAIDTCRHYLAIVKKVCRKAYKEGHAD; 28 residues) was a predicted NonHemo, putatively novel candidate assigned to Tier1_A. It had a mean pLDDT of 91.68 and passed the available short water-phase MD quality-control assessment with a mean RMSD of 2.73 Å. GEAMP_446036444fc498a1 (eAMP-02; GAMEKAKKVRQRCGEVFRYAIVTGRAIYN; 29 residues) was also a predicted NonHemo, putatively novel Tier1_A candidate, with a mean pLDDT of 89.67 and strong dual-model AMP support. The two candidates were therefore retained as independent safety-oriented candidates, despite their cysteine content requiring attention during later synthesis and oxidation-state control.

**Table 1.**
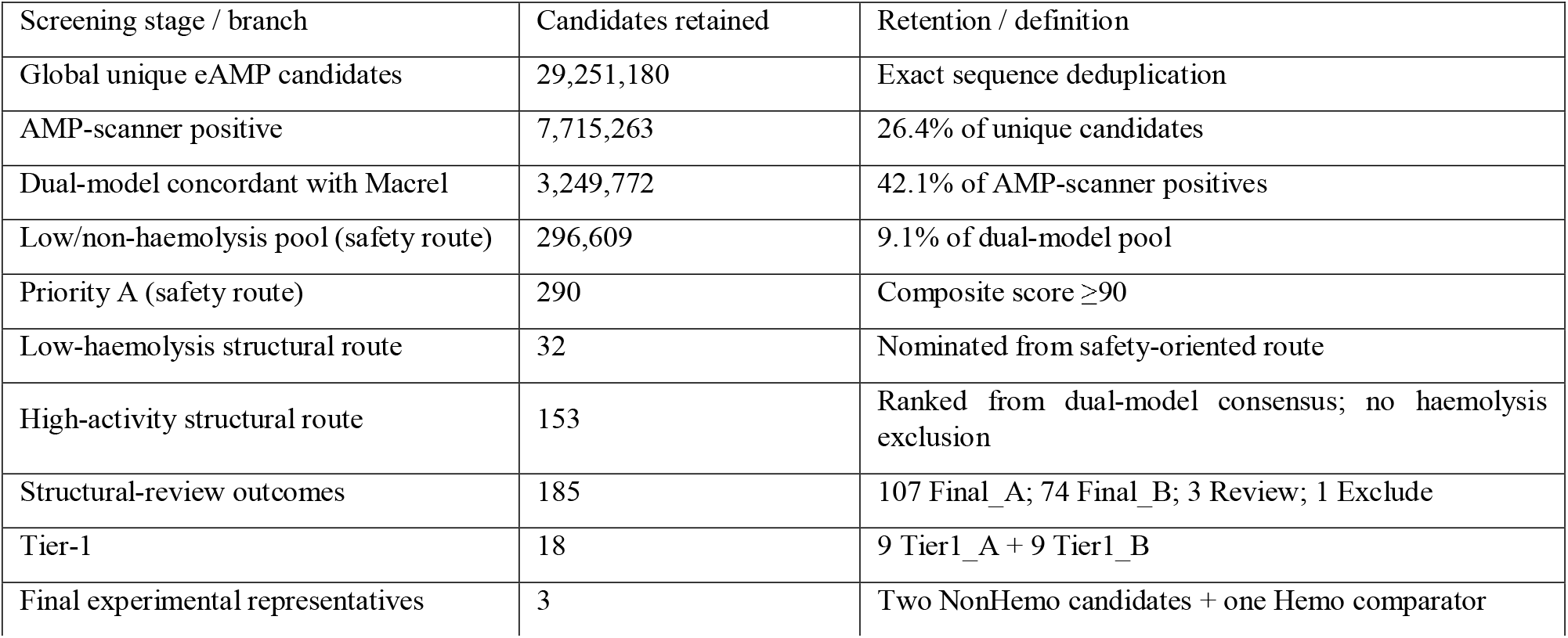
Audit trail of candidate filtering and branch-specific structural nomination.

| Screening stage / branch | Candidates retained | Retention / definition |
| --- | --- | --- |
| Global unique eAMP candidates | 29,251,180 | Exact sequence deduplication |
| AMP-scanner positive | 7,715,263 | 26.4% of unique candidates |
| Dual-model concordant with Macrel | 3,249,772 | 42.1% of AMP-scanner positives |
| Low/non-haemolysis pool (safety route) | 296,609 | 9.1% of dual-model pool |
| Priority A (safety route) | 290 | Composite score $\geq 90$ |
| Low-haemolysis structural route | 32 | Nominated from safety-oriented route |
| High-activity structural route | 153 | Ranked from dual-model consensus; no haemolysis exclusion |
| Structural-review outcomes | 185 | 107 Final_A; 74 Final_B; 3 Review; 1 Exclude |
| Tier-1 | 18 | 9 Tier1_A + 9 Tier1_B |
| Final experimental representatives | 3 | Two NonHemo candidates + one Hemo comparator |

**Table 2.** Structural-review outcomes for the 185 candidates nominated by the two prioritisation routes.

| Route | Purpose | Input | Final_A | Final_B | Other |
| --- | --- | --- | --- | --- | --- |
| Low-haemolysis route | Selectivity-oriented candidates | 32 | 15 | 13 | 3 Review; 1 Exclude |
| High-activity route | Membrane-active comparator route | 153 | 92 | 61 | 0 Review; 0 Exclude |
| Total | Parallel structural review | 185 | 107 | 74 | 3 Review; 1 Exclude |

**Table 3.** Final sequence-defined eAMP representatives and experimental roles.

| Candidate | Sequence | Class | Key evidence | pLDDT | Experimental role |
| --- | --- | --- | --- | --- | --- |
| eAMP-01<br>GEAMP_71c13939<br>3ac596b5 | GLAIDTCRHYLAIVKKV<br>CRKAYKEGHAD | NonHemo<br>putatively<br>novel | Tier1_A; water MD<br>pass; RMSD 2.73 Å | 91.68 | Primary safety<br>candidate; 2 Cys |
| eAMP-02<br>GEAMP_44603644<br>4fc498a1 | GAMEKAKKVRQRCGE<br>VFRYAIVTGRAIYN | NonHemo<br>putatively<br>novel | Tier1_A; strong dual-model support | 89.67 | Independent<br>safety candidate;<br>1 Cys |
| eAMP-03<br>GEAMP_12ffb5d58<br>9c8cb1b | VVKRYIKSIGKGILKVM<br>SKMGI | Hemo<br>high-activity | Final_A; deep<br>insertion; POPG=0.292 | 83.75 | Membrane/selecti-<br>vity comparator |

**Table 4.** Concentration-dependent colony-count response of the three final eAMP candidates against *E. coli*, expressed relative to the matched 8 μM condition.

| Candidate | 8 $\mu\text{M}$ | 16 $\mu\text{M}$ | 32 $\mu\text{M}$ | 64 $\mu\text{M}$ | 128 $\mu\text{M}$ | 128 $\mu\text{M}$ relative<br>to matched 8<br>$\mu\text{M}$ |
| --- | --- | --- | --- | --- | --- | --- |
| eAMP-01 | 1,754 | 1,315 | 814 | 417 | 174 | 90.1% lower;<br>1.00-log<sub>10</sub> vs 8<br>$\mu\text{M}$ |
| eAMP-02 | 1,662 | 1,119 | 618 | 306 | 131 | 92.1% lower;<br>1.10-log10 vs 8<br>μM |
| eAMP-03 | 1,601 | 741 | 316 | 179 | 48 | 97.0% lower;<br>1.52-log10 vs 8<br>μM |

**Table 5.** Concentration-dependent colony-count response of the three final eAMP candidates against *S. aureus*, expressed relative to the matched 8 μM condition.

| Candidate | 8 μM | 16 μM | 32 μM | 64 μM | 128 μM | 128 μM relative<br>to matched 8<br>μM |
| --- | --- | --- | --- | --- | --- | --- |
| eAMP-01 | 3,621 | 2,282 | 1,489 | 699 | 215 | 94.1% lower;<br>1.23-log10 vs 8<br>μM |
| eAMP-02 | 3,798 | 2,189 | 1,443 | 645 | 176 | 95.4% lower;<br>1.33-log10 vs 8<br>μM |
| eAMP-03 | 3,574 | 1,491 | 646 | 191 | 19 | 99.5% lower;<br>2.27-log10 vs 8<br>μM |

**Figure 2.**
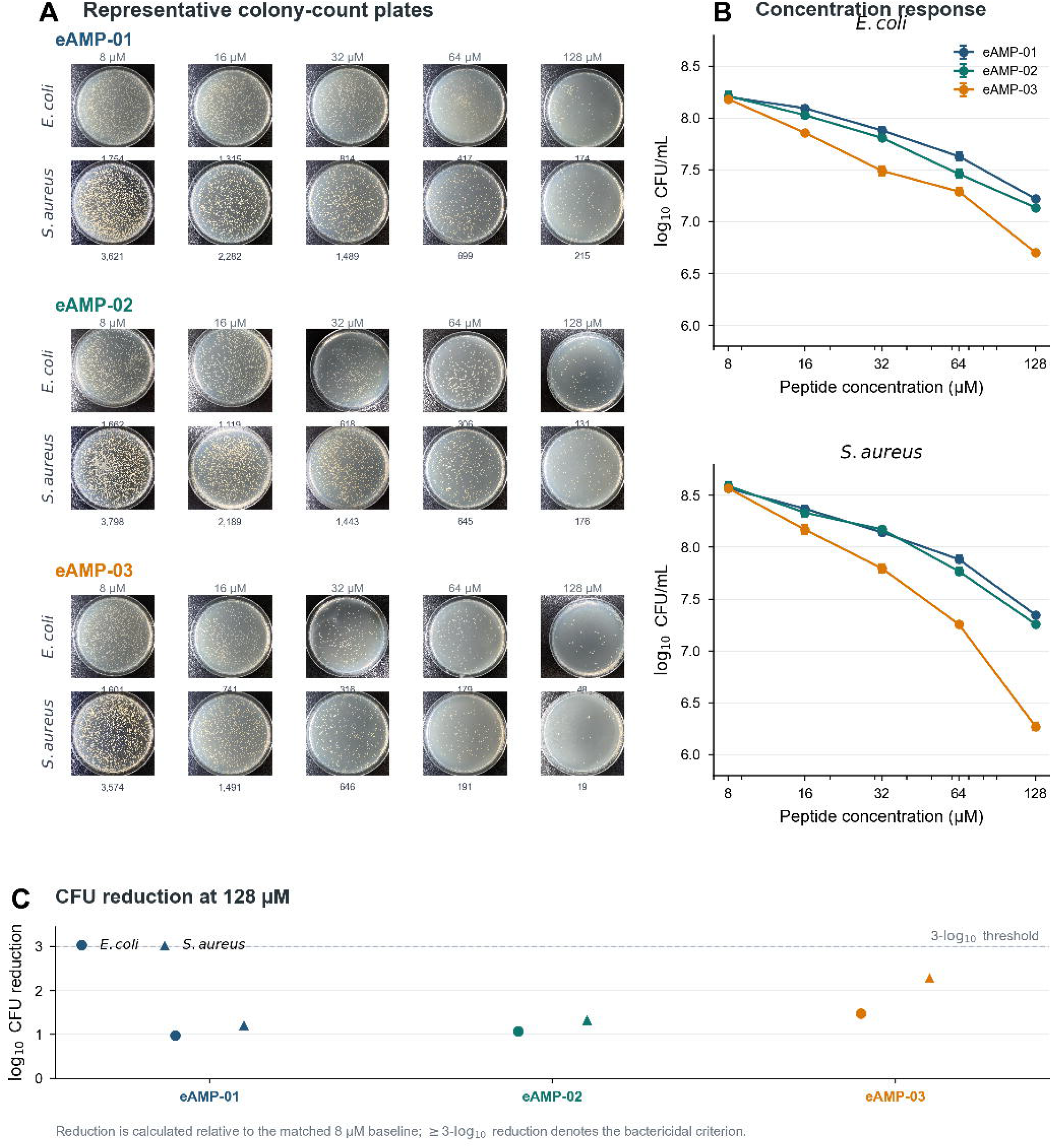
Structural prioritisation of the three final eAMP candidates. GEAMP_71c139393ac596b5 and GEAMP_446036444fc498a1 represent predicted NonHemo, safety-oriented Tier1_A candidates, whereas GEAMP_12ffb5d589c8cb1b represents a predicted Hemo, high-activity comparator. Structural panels summarize sequence length, net charge, mean pLDDT, predicted α-helical fraction, and relevant developability annotations.

GEAMP_12ffb5d589c8cb1b (eAMP-03; VVKRYIKSIGKGILKVMSKMGI; 22 residues) was deliberately retained from the high-activity route as a predicted haemolytic comparator rather than a therapeutic candidate. It met Final_A structural criteria, carried strong dual-model AMP support, and showed the strongest available mechanism-oriented membrane evidence. In the MARTINI membrane simulation, the peptide reached an interface/deep-insertion state with a POPG contact fraction of 0.292. This result supports a testable membrane-association hypothesis but does not by itself establish membrane disruption as the dominant antibacterial mechanism.

#### Candidate-specific MD supported stability and membrane-association hypotheses

All three candidates (eAMP-01, eAMP-02, and eAMP-03) were subjected to molecular-dynamics (MD) simulations in both aqueous and membrane environments (Fig. 3). Water-phase stability (Fig. 3A, 5-ns MD): the backbone RMSD and per-residue RMSF of the three candidates were evaluated during the 5-ns simulation. The backbone RMSD remained low (≤6 Å) for all three candidates, and the RMSF showed a normal increase only at the terminal residues while remaining low in the helical core, supporting relatively stable conformations during the short simulation.

**Figure 3.**
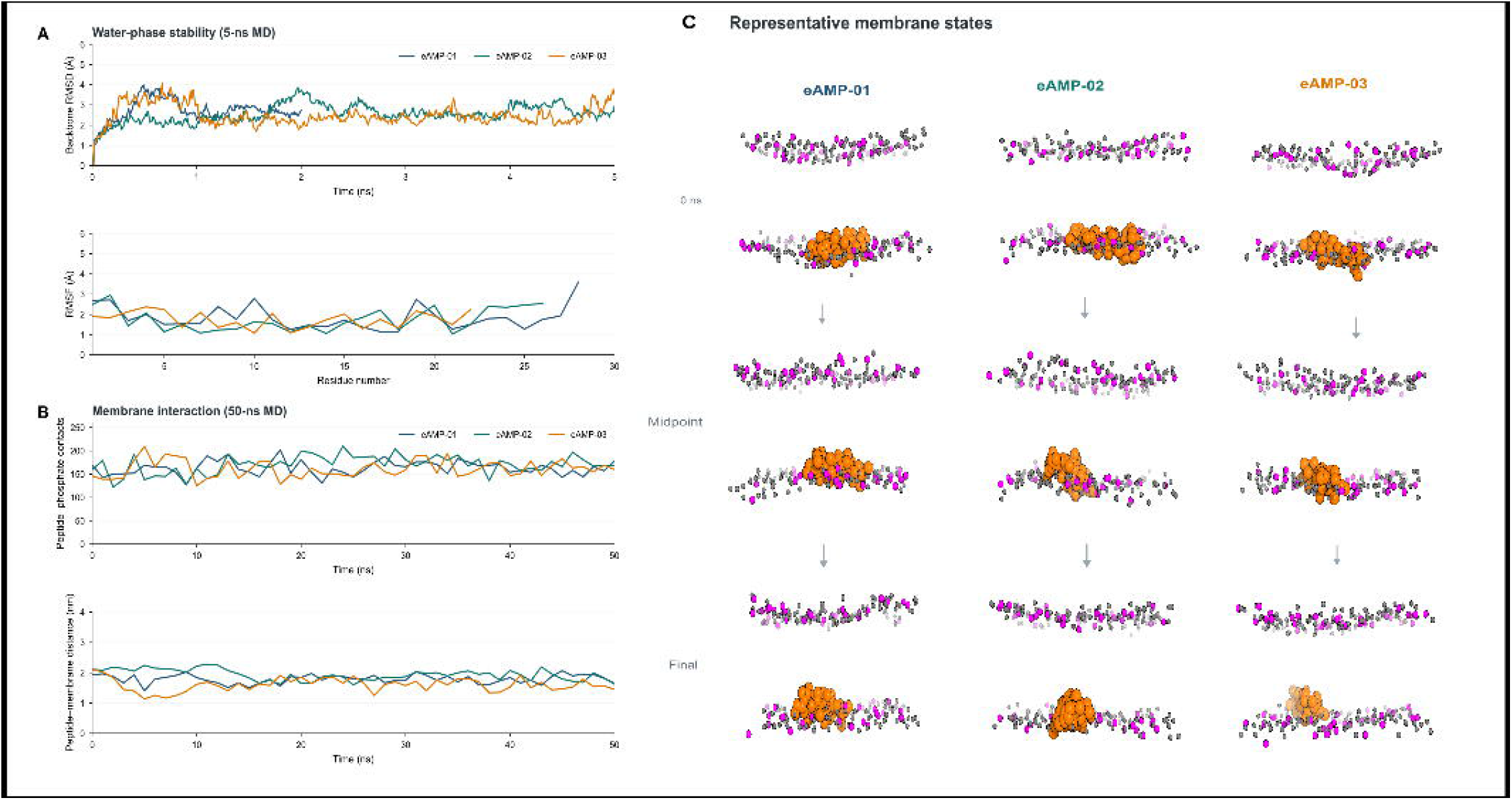
Molecular-dynamics evidence for the three final eAMP candidates. (A) Water-phase stability over 5 ns, showing backbone RMSD and per-residue RMSF for eAMP-01, eAMP-02, and eAMP-03. (B) Membrane interaction over 50 ns, showing peptide-phosphate contacts and peptide-membrane distance for all three candidates. (C) Representative membrane states at 0 ns, midpoint, and final time points. These are single pilot trajectories that support descriptive comparisons but not statistical inference or quantitative mechanistic ranking.

Membrane interaction (Fig. 3B, 50-ns MD): the peptide-phosphate contact number and peptide-membrane distance of the three candidates were tracked during the 50-ns membrane simulation. The peptide-phosphate contact number increased and tended to stabilise, while the peptide-membrane distance decreased and remained near the membrane interface, indicating that all three candidates showed a continuous tendency to associate with the anionic model membrane. Representative membrane states (Fig. 3C): snapshots at 0 ns, midpoint, and final time points showed that all three candidates gradually approached the membrane interface from an initial solution or surface state and ultimately adopted surface-bound or insertion-like conformations. These single pilot trajectories collectively support conformational-stability and membrane-association hypotheses for the three candidates; membrane disruption remains to be verified by membrane-permeability, depolarisation, or liposome-leakage assays.

#### Replicated colony-count assays showed concentration-dependent relative reductions (Fig. 4)

The same three sequence-defined GEAMP candidates were tested against E. coli and *S. aureus* at 8, 16, 32, 64, and 128 μM using three recorded replicate plate series with linked raw images. Across both organisms, colony counts decreased monotonically from 8 to 128 μM for all three peptides. Because the experimental records did not include an untreated or time-zero CFU reference, the 8 μM condition was used only as a within-candidate, within-organism reference for describing relative changes across the tested concentration series.

**Figure 4.**
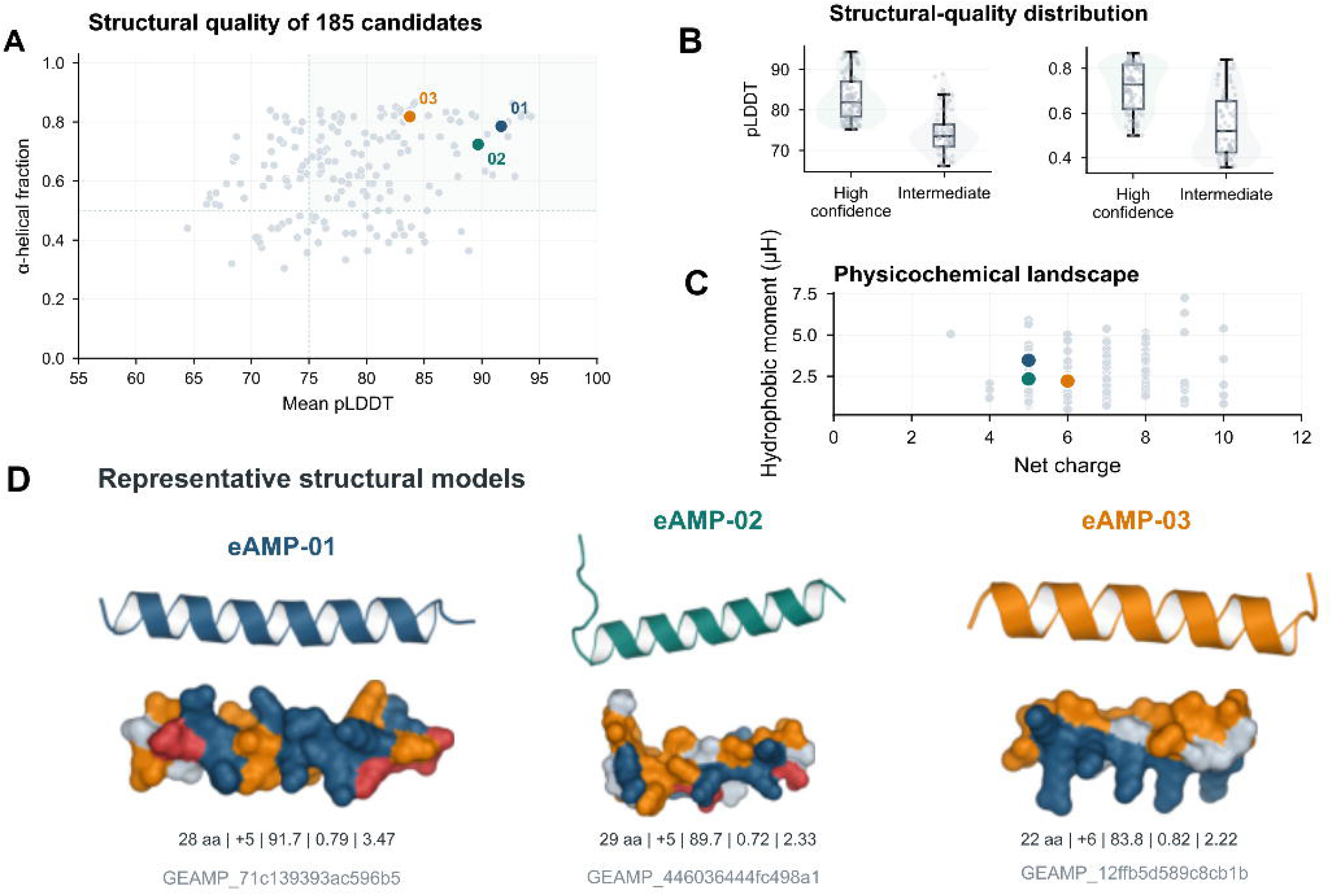
Replicated colony-count profiles of the three sequence-matched final eAMP candidates. Representative plates and quantitative concentration-response summaries are shown for GEAMP_71c139393ac596b5 (eAMP-01), GEAMP_446036444fc498a1 (eAMP-02), and GEAMP_12ffb5d589c8cb1b (eAMP-03) against E. coli and *S. aureus* across 8–128 μM. Rep1–Rep3 correspond to three recorded plate series with linked raw images. All three peptides showed monotonic colony reduction with increasing concentration. Relative log10 reductions were calculated against the matched 8 μM condition because an untreated or time-zero CFU reference was not available in the source records. These within-series relative reductions are not formal MBC measurements.

Against *E. coli*, eAMP-01 decreased from 1,754 colonies at 8 μM to 174 at 128 μM, corresponding to a 90.1% lower colony count and a 1.00-log10 relative reduction versus the matched 8 μM condition. eAMP-02 decreased from 1,662 to 131 colonies, corresponding to a 92.1% decrease and a 1.10-log10 relative reduction. eAMP-03 showed the largest change, decreasing from 1,601 colonies at 8 μM to 48 at 128 μM, corresponding to a 97.0% decrease and a 1.52-log10 relative reduction.

The same endpoint ordering was observed against *S. aureus*. eAMP-01 decreased from 3,621 colonies at 8 μM to 215 at 128 μM, corresponding to a 94.1% lower colony count and a 1.23-log10 relative reduction versus the matched 8 μM condition. eAMP-02 decreased from 3,798 to 176 colonies, corresponding to a 95.4% decrease and a 1.33-log10 relative reduction. eAMP-03 again showed the largest endpoint change, decreasing from 3,574 colonies to 19 at 128 μM, corresponding to a 99.5% decrease and a 2.27-log10 relative reduction.

The endpoint ordering was therefore eAMP-03 > eAMP-02 > eAMP-01 in both organisms. The comparatively strong concentration-dependent response of eAMP-03 is directionally consistent with its strong dual-model AMP support and deep membrane-insertion simulation phenotype, but its predicted Hemo classification precludes interpretation as the preferred therapeutic candidate without direct haemolysis and cytotoxicity measurements. Among the two predicted NonHemo candidates, eAMP-02 showed the larger relative endpoint reduction in both organisms. These measurements quantify changes relative to the matched 8 μM condition; because no untreated or time-zero CFU reference was available, they do not support calculation of a formal MBC.

## Discussion

The principal contribution of this work is an auditable prioritisation architecture for converting bacterial proteome space into experimentally testable encrypted AMP candidates. The workflow does not rely on one classifier or one extraction rule. Instead, it combines three layer-specific extraction scores, dual-model AMP prediction, two complementary downstream prioritisation routes, structure-aware review, focused developability and novelty assessment, and candidate-specific MD evidence. The numerical audit trail—from 29.25 million unique peptide sequences to a 3.25-million-member dual-model consensus, 185 structurally reviewed candidates, 18 Tier-1 candidates, and three final representatives—makes the selection process inspectable while preserving route-specific provenance.

A key design feature is the branch-specific treatment of predicted haemolysis. The low-haemolysis route applied NonHemo/safety-oriented filtering to nominate selectivity-oriented candidates, whereas the high-activity route ranked the same dual-model consensus without excluding predicted haemolytic sequences and retained haemolysis status as a risk annotation. The lack of overlap between the two top-200 ranking sets further indicates that these routes captured distinct prioritisation objectives. GEAMP_71c139393ac596b5 and GEAMP_446036444fc498a1 emerged from the safety-oriented route, whereas GEAMP_12ffb5d589c8cb1b was deliberately retained as a high-activity comparator.

The sequence-level design carries the same GEAMP identifiers and peptide sequences into the colony-count assay. All three candidates showed monotonic concentration-dependent colony reduction in both organisms, with the same endpoint ordering of eAMP-03 > eAMP-02 > eAMP-01 at 128 μM. Relative to the matched 8 μM condition, eAMP-03 showed the largest decrease, particularly against *S. aureus* (99.5% lower colony count; 2.27-log10 relative reduction), which is directionally consistent with its strong computational AMP support and deep membrane-insertion phenotype. However, the same candidate is predicted Hemo, illustrating the potential trade-off between antibacterial activity and host-cell selectivity that motivated the two-route design.

The two predicted NonHemo candidates showed smaller but still monotonic concentration-dependent changes. eAMP-02 exceeded eAMP-01 at the 128 μM endpoint in both E. coli (1.10 vs 1.00 log10 relative reduction) and *S. aureus* (1.33 vs 1.23 log10 relative reduction), making eAMP-02 the higher-priority safety-oriented candidate for direct haemolysis and cytotoxicity testing. These values should not be interpreted as growth-control-based inhibition or bactericidal log reductions: the experimental records lacked an untreated or time-zero CFU reference, and the calculations use the matched 8 μM condition as an internal baseline. A formal MBC therefore cannot be determined from the current dataset.

For the MD component, all three representatives now have candidate-specific water-phase (5 ns) and membrane-environment (50 ns) evidence; these single pilot trajectories support descriptive comparisons of conformational stability and membrane association, but, without independent replicate simulations, they do not support statistical inference or quantitative mechanistic ranking.

Future validation should prioritise experimental haemolysis and cytotoxicity, standardised biological repeat assays, time-kill kinetics, and direct membrane-function measurements such as depolarisation, permeabilisation, or vesicle-leakage assays. Parent-protein processing experiments would be particularly valuable because they would test the defining biological premise of an encrypted peptide: that an active fragment can actually be generated from its larger precursor. These experiments would transform the current discovery framework from a sequence- and mechanism-informed prioritisation system into a more complete eAMP biology study.

## Conclusions

This study establishes a traceable multi-layer route from bacterial nr95 proteomes to experimentally evaluated encrypted AMP candidates. Three complementary extraction modes with layer-specific scoring generated 29.25 million unique peptide hypotheses, which were reduced by dual-model AMP prediction and then prioritised through independent low-haemolysis and high-activity routes before structure prediction, structural review, Tier-1 assessment, novelty analysis, and candidate-specific MD. Three sequence-defined representatives were advanced to replicated colony-count testing against E. coli and *S. aureus* and showed monotonic concentration-dependent reductions across 8–128 μM. eAMP-03 showed the largest relative endpoint decrease in both organisms, whereas eAMP-02 showed the larger response among the two predicted NonHemo candidates. Because the available experimental records use the matched 8 μM condition as an internal reference and do not contain an untreated or time-zero CFU baseline, these data support relative concentration-response comparisons but not formal MBC assignment. Experimental haemolysis, cytotoxicity, controlled bactericidal assays, and direct evidence of peptide release from parent proteins remain necessary before therapeutic-lead or endogenous-eAMP claims are made.

## Materials and Methods

### Proteome input and preprocessing

The discovery input comprised nr95 non-redundant protein collections from 265 high-quality bacterial genomes. Each protein set was generated by redundancy reduction at 95% sequence identity, and only parent proteins of at least 100 amino acids were retained for eAMP mining. A fixed input list was used throughout the analysis to preserve genome-level and parent-protein provenance.

### Layer-specific extraction scoring

The three extraction layers used distinct eight-point scoring functions. For L1 terminal candidates, the relative peptide score (RPS; maximum 8) comprised terminal proximity, peptide length, cleavage context, and net positive charge, each contributing up to 2 points. Terminal proximity scored 2 when a fragment began within 60 residues of the N terminus or ended within 60 residues of the C terminus and 1 when the corresponding distance was ≤100 residues. Length scored 2 for 18–35 residues and 1 for 12–17 or 36–40 residues. Cleavage context scored 2 for a basic motif or K/R flank and 1 for adjacent G/A/S/P. Net charge scored 2 for charge ≥+3 and 1 for charge +1 to +2. L1 candidates were retained at RPS ≥7. L2 and L3 did not use this terminal RPS. The L2 score comprised cleavage_score_v2 (basic motif=4, K/R flank=3, adjacent G/A/S/P=2), length_score_v2 (22–30 residues=2), and charge_score_v2 (net charge ≥+5=2), with a maximum of 8 and a retention threshold of 8. The L3 score comprised charge_score_v3 (≥+7=4, ≥+5=3, ≥+4=2, ≥+3=1), length_score_v3 (22–30 residues=2), and hydrophobic_score_v3 (hydrophobic fraction 0.35–0.50=2), again with a maximum of 8 and a retention threshold of 8.

### Three-layer eAMP candidate extraction

Layer 1 (L1) scanned the N- and C-terminal 60-residue regions of eligible parent proteins using peptide windows of 20–32 residues and retained fragments meeting the L1 RPS threshold (≥7). Layer 2 (L2) identified K, R, KR, RR, RK, and KK cleavage-like motifs and examined ±20-residue neighbourhoods around these sites, retaining only candidates that achieved the maximum L2 score of 8. Layer 3 (L3) first identified 30-residue internal windows with net charge ≥+4, designated these regions as cationic hotspots, and then performed fine-scale peptide extraction within each hotspot, retaining candidates with an L3 score of 8. L1, L2, and L3 generated 12,070,288, 8,111,517, and 12,483,768 sequence records, respectively. Results were merged and globally deduplicated by exact amino-acid sequence to yield 29,251,180 unique candidates; multi-layer origin was retained as provenance metadata.

### Dual-model AMP prediction

AMP-scanner v2 was run under Python 3.7 and TensorFlow 1.15. Candidates with an AMP-scanner probability >0.5 were considered positive, yielding 7,715,263 candidates. These sequences were subsequently analysed with Macrel v1.6.0, and only sequences classified as positive by both models were retained. The dual-model consensus contained 3,249,772 candidates. Agreement between the two models was used to reduce dependence on a single prediction framework rather than as independent experimental confirmation of antimicrobial activity [6,7].

### Safety-oriented haemolysis filtering and 100-point composite prioritisation

For the safety-oriented route, candidates classified as NonHemo by Macrel or assigned a predicted haemolysis probability <0.5 by the haemolysis-prediction layer were retained, reducing the 3,249,772-member dual-model consensus to 296,609 sequences. These candidates were scored on a 100-point scale comprising AMP-related physicochemical features (maximum 40 points), model evidence (30 points), cross-species conservation (20 points), and safety evidence (10 points). The physicochemical component prioritised peptide length in the 12–40-residue range, net positive charge, and a hydrophobic fraction centred on 0.30–0.55. The model component integrated AMP-scanner and Macrel evidence, the conservation component rewarded cross-species recurrence, and the safety component rewarded NonHemo status while applying a haemolysis penalty where relevant. Priority A was defined as score ≥90, Priority B as 80–89, Priority C as 70–79, and Priority D as <70; the corresponding candidate counts were 290, 177,457, 115,447, and 3,415. This scoring scheme applied to the safety-oriented route and was not used to exclude predicted haemolytic candidates from the independent high-activity route.

### Branch-specific structural candidate nomination and review

Structural candidate nomination was performed through two parallel routes from the 3,249,772-member dual-model consensus. The low-haemolysis route applied the safety-oriented filtering and prioritisation described above and nominated 32 candidates for structure prediction. The high-activity route did not apply haemolysis exclusion; predicted haemolysis was retained as a risk annotation, and candidates were ranked by the predefined HighActivityScore, yielding 153 structural candidates. The top-200 ranking sets from the two routes showed 0% overlap. OmegaFold and ColabFold were then used to generate structural models for all 185 nominated candidates. Structural review combined batch image quality control, functional-class quality control, mean pLDDT, predicted α-helical fraction, source priority, and manual three-dimensional inspection in PyMOL. Final_A required mean pLDDT ≥75, predicted α-helical fraction ≥0.50, image-QC pass, and Priority A source status; Final_B required mean pLDDT ≥65 and predicted α-helical fraction ≥0.35. The low-haemolysis route yielded 15 Final_A, 13 Final_B, three Review, and one Exclude candidates; the high-activity route yielded 92 Final_A and 61 Final_B candidates. Overall, 107 candidates were classified Final_A, 74 Final_B, three Review, and one Exclude.

### Tier-1 developability assessment

Eighteen candidates entered Tier-1 assessment. The assessment included sequence-liability checks, physicochemical and aggregation-related properties, synthesis feasibility, membrane-activity prediction, available short water-phase MD evidence, and novelty assessment. Sequence review identified 17 low-risk candidates and one medium-risk candidate, with no high-risk candidates; 11 candidate pairs showed >85% sequence similarity. No candidate was assigned high aggregation risk. Synthesis feasibility was classified as easy for 17 candidates and moderate for one. Membrane-activity prediction classified 12 candidates as strong and six as moderate. The final set was divided into nine Tier1_A and nine Tier1_B candidates.

### Candidate annotation and novelty analysis

Candidate annotation integrated source and functional information with downstream validation annotations. Novelty assessment combined database-similarity searches, sequence clustering, and motif inspection. These results were used to distinguish putatively novel, potentially novel, and previously similar sequence classes for prioritisation. Novelty labels were not interpreted as direct evidence of a previously unrecognised biological family without independent functional and evolutionary validation.

### Water-phase and membrane molecular-dynamics analysis

For the three final representatives, candidate-specific short water-phase (5 ns) simulations provided backbone RMSD and per-residue RMSF traces; these were used to assess conformational stability across the short simulation.

Membrane-environment pilot simulations (50 ns) used a MARTINI coarse-grained representation and, for all three final representatives, evaluated peptide-phosphate contacts, peptide-membrane distance, and representative membrane-state snapshots at 0 ns, midpoint, and final time points. GEAMP_12ffb5d589c8cb1b reached an interface/deep-insertion state with a POPG contact fraction of 0.292.

### Final candidate selection

Three candidates were selected to represent complementary experimental hypotheses. GEAMP_71c139393ac596b5 (GLAIDTCRHYLAIVKKVCRKAYKEGHAD) was a predicted NonHemo, putatively novel Tier1_A candidate with mean pLDDT 91.68 and passing short water-phase MD evidence; it was designated the primary safety-oriented candidate. GEAMP_446036444fc498a1 (GAMEKAKKVRQRCGEVFRYAIVTGRAIYN) was a predicted NonHemo, putatively novel Tier1_A candidate with mean pLDDT 89.67 and strong dual-model support and was retained as an independent safety-oriented candidate. GEAMP_12ffb5d589c8cb1b (VVKRYIKSIGKGILKVMSKMGI) was a predicted Hemo, high-activity Final_A candidate with strong dual-model support and deep membrane-insertion evidence; it was retained as a high-activity mechanism and selectivity comparator.

#### Replicated colony-count assay and relative reduction analysis

The three final sequence-defined candidates were tested against E. coli and *S. aureus* at 8, 16, 32, 64, and 128 μM in twofold concentration steps. Rep1, Rep2, and Rep3 were recorded as three replicate plate series, with each replicate linked to its corresponding raw plate image. Raw colony counts and derived CFU/mL values were recorded for each candidate-organism-concentration combination.

CFU/mL was calculated as colony count divided by the product of dilution fraction and plated volume in millilitres: CFU/mL = colony count / (dilution fraction × plated volume [mL]). The available source records did not include an untreated growth-control or time-zero CFU value suitable for bactericidal log-reduction analysis. Accordingly, the matched 8 μM condition was used as an internal reference within each peptide-organism series, and relative log10 reduction at concentration C was calculated as log10[CFU/mL at 8 μM ÷ CFU/mL at C], with the 8 μM condition defined as 0. These values describe concentration-dependent changes relative to the lowest tested peptide concentration and are not equivalent to growth-control-based inhibition or formal bactericidal log reduction. No MBC was assigned from this dataset.

## Data Availability

The datasets generated and analysed during this study, including candidate catalogues, prediction and scoring outputs, structural-review records, Tier-1 assessments, novelty annotations, MD summaries, raw colony counts, calculated CFU values, relative log10-reduction tables, and replicate plate images, are available from the corresponding author on reasonable request. The associated genome accession list, parent-protein provenance records, and final peptide-sequence registry are maintained with the study data package.

## Code Availability

Analysis scripts used for three-layer extraction, layer-specific scoring, dual-model prediction, branch-specific prioritisation, structural review, Tier-1 assessment, novelty annotation, MD analysis, and figure and supplementary-table generation, together with the corresponding environment specifications, are available from the corresponding author on reasonable request.

## Author Contributions

Qingxiu Li: methodology, computational analysis, experimental work, data curation, and manuscript drafting. Zhenjun Li: conceptualization, supervision, manuscript review, and editing.

## Competing Interests

The authors declare no competing interests.

## Funding

This research received no specific grant from any funding agency in the public, commercial, or not-for-profit sectors.

## Ethics Approval

No human participants or vertebrate animals were involved in this study; institutional ethics approval was not required.

## Supplementary Tables

Table S1, integrated annotation catalogue for the 185 structurally reviewed candidates; Table S2, physicochemical and selection parameters for the three final candidates; Table S3, water-phase and membrane molecular-dynamics evidence; Table S4, MIC results for the 20-candidate screen; Table S5, three-replicate MBC colony counts; and Table S6, data-source and definition notes.

## Notes

### Competing Interest Statement

The authors have declared no competing interest.

